# Pathways analyses of schizophrenia GWAS focusing on known and novel drug targets

**DOI:** 10.1101/091264

**Authors:** H.A. Gaspar, G. Breen

## Abstract

Using successful genome-wide association results in psychiatry for drug repurposing is an ongoing challenge. Databases collecting drug targets and gene annotations are growing and can be harnessed to shed a new light on psychiatric disorders. We used GWAS summary statistics from the Psychiatric Genetics Consortium (PGC) Schizophrenia working group and a repositioning model for schizophrenia by testing the enrichment of antipsychotics. As sample size increases, schizophrenia GWAS results show increasing enrichment for known antipsychotic drugs, selective calcium channel blockers, and antiepileptics. Each of these therapeutical classes targets different gene subnetworks. We identify 162 Bonferroni-significant druggable genes, and 128 FDR-significant biological pathways related to neurons, synapses, genic intolerance, membrane transport, epilepsy, and mental disorders. These results suggest that, in schizophrenia, current well-powered GWAS results can reliably detect known schizophrenia drugs and thus may hold considerable potential for the identification of new therapeutic leads. Moreover, antiepileptics and calcium channel blockers may provide repurposing opportunities. This study also reveals significant pathways in schizophrenia that were not identified previously, and provides a workflow for pathway analysis and drug repurposing using GWAS results.

## INTRODUCTION

Genome-wide association studies (GWAS) have been performed on numerous human disorders and traits ^1^, uncovering thousands of associations between disorders or quantitative phenotypes and common genetic variants, usually single nucleotide polymorphisms (SNPs), that ‘tag’ or identify specific genetic loci. Summary statistics from hundreds of GWASs are freely available online, including those from the Psychiatric Genetics Consortium (PGC) Schizophrenia working group. Schizophrenia is a complex disorder with a lifetime prevalence of ∼1%, significant environmental risk factors, and a heritability of 65%-85%^2^ that has been suggested to be highly polygenic in nature ^3^. As with other complex genetic disorders, the application of GWAS to schizophrenia has identified multiple disease susceptibility loci. In 2014, over 100 robustly associated loci were identified in a GWAS meta-analysis by the PGC ^4^. Similar progress is underway in other psychiatric disorders, with new GWAS reports expected for attention deficit hyperactivity disorder, autism, major depressive disorder, anorexia nervosa, and bipolar disorder in the next year. However, a key question arises: how can the emergence of new and well powered GWAS data inform the development of new therapeutics?

Most attention on the therapeutic utility of GWAS has focused on the identification of individual drug targets ^5^. Nelson et al. recently demonstrated that the proportion of drug mechanisms with genetic support increases from 2.0% at the preclinical stage to 8.2% after successful approval ^6^. Results from genetic studies can also guide repurposing - the finding of new indications for known drugs ^7–9^. Recent studies have also shown how pathway analysis on GWAS data could help discover new drugs for schizophrenia ^10,11^. However, these studies, as well as studies focused on single genes or targets, have generally lacked a validation step to show if a GWAS has sufficient power to reliably identify known drugs: a crucial indication that would lend confidence to the discovery of novel drug associations in GWAS data.

Mining of data available on drug-gene interactions (**Fig. 1**) allows the combination of individual drug targets into “drug pathways” represented by sets of genes that encode all targets of a given drug or potential novel therapeutic. Any drug can be represented by such a gene-set derived from its drug activity profile, and assigned a p-value generated by pathway analysis assessing the association of a given drug gene-set with the phenotype. An enrichment curve can be drawn for any particular group of drugs using the entire dataset of drugs ranked by p-value. The associated area under the enrichment curve (AUC) provides a simple way to assess the enrichment of any class of drug for a specific disorder. To validate a drug repurposing model, we propose to test the enrichment of a known class of drugs: antipsychotics for schizophrenia, anxiolytics for anxiety disorders, etc.

**Figure 1.**
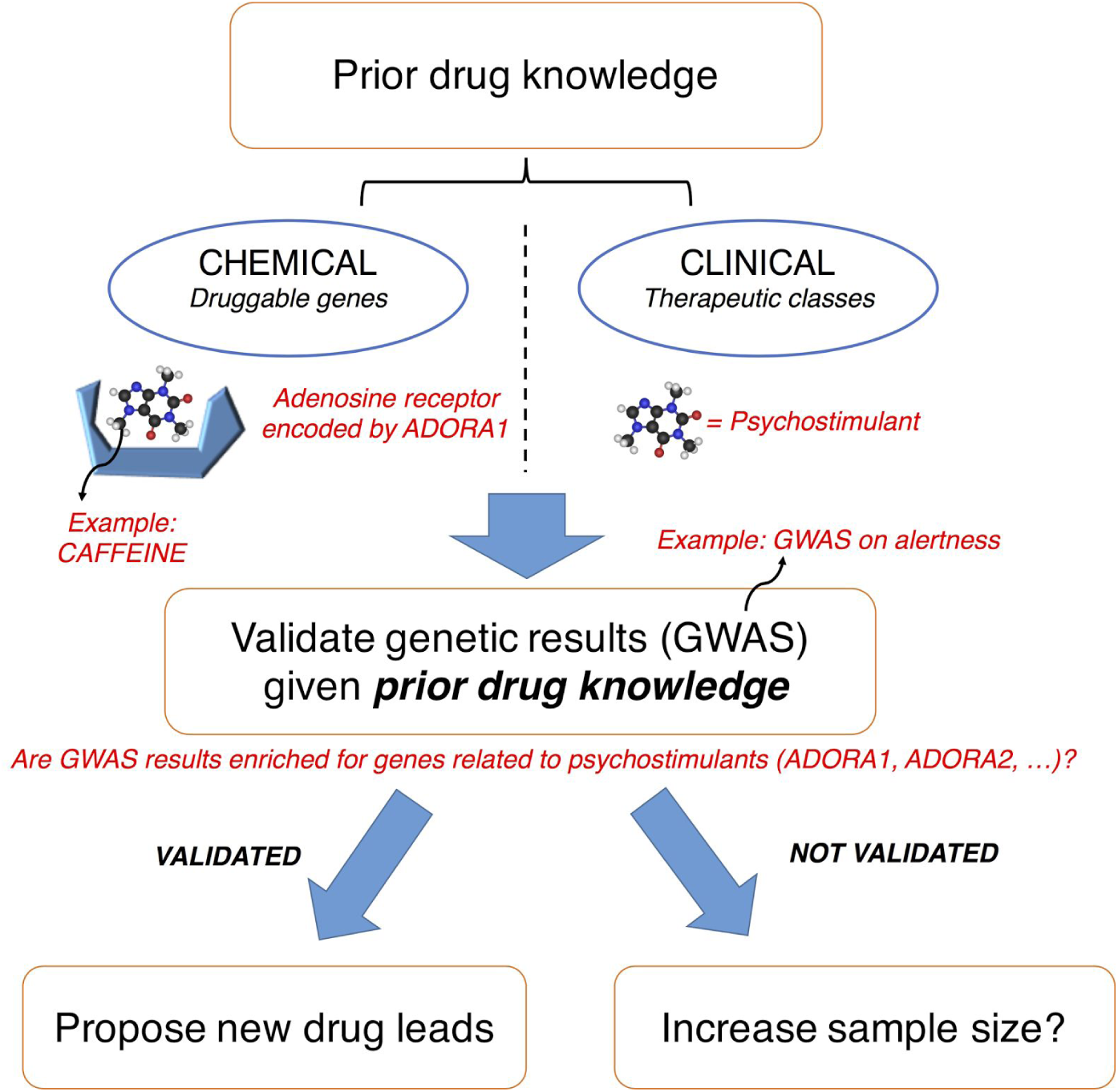
Using drug knowledge to validate genetic results. Drug knowledge, encompassing therapeutic classes and druggable genes (*e.g*., caffeine is a psychostimulant targeting adenosine receptors), may be used to validate the ability of a GWAS to find known drugs for a given trait (*e.g.*, alertness). Novel targets and potential drugs could then be found in validated genetic results.

In this article, we performed pathway analysis to assess the significance of drugs in schizophrenia GWAS. We analysed and compared three successively larger schizophrenia studies from the PGC Schizophrenia working group: SCZ-PGC1 ^12^, SCZ-PGC1+SWE ^13^, and SCZ-PGC2 ^4^. We also analysed the complete SCZ-PGC2 GWAS for the associations of gene families, gene ontology (GO) pathways, canonical pathways, disease pathways, drugs and drug classes with schizophrenia. However, a common problem in pathway analysis is the interpretation of the top pathways. We propose a new workflow to visualise and cluster significant biological pathways by accounting for pathway similarities as well as pathway significance, based on a kernel variant of the Generative Topographic Mapping approach^14,15^.

## RESULTS

### Druggable genes

An analysis of druggable genes was conducted using SCZ-PGC2. A druggable gene Manhattan plot is presented in **Fig. S1a in Supplement 1**. We define druggable genes as genes with known drug interactions and genetic variations. After applying a Bonferroni correction, 162 out of 3053 are significant for schizophrenia, and 505 have a Benjamini and Hochberg FDR q-value < 5%. Among significant genes, several are related to the major histocompatibility complex (MHC), calcium voltage-gated channels, potassium channels or cholinergic receptors (**Fig. S1b in Supplement 1**). Among cholinergic genes, the cluster of genes *CHRNA3-CHRNA5-CHRNB4* is strongly associated with schizophrenia, whereas *CHRNA4* is FDR-significant but not Bonferroni-significant; *CHRM4* is Bonferroni-significant. Three CACN genes show a Bonferroni-significant association with schizophrenia: *CACNA1I*, *CACNA1C* and *CACNB2*. *DRD2* and *HTR5A* are the antipsychotic target genes with the strongest association amongst the dopamine and 5-HT receptors families.

### Pathway Maps

We built pathway maps colored by association with SCZ-PGC2 using the kernel generative topographic mapping approach (k-GTM) to identify significant gene-sets with similar gene content. The top 50 biological pathways from SCZ-PGC2 pathway analysis were thus mapped onto a 2D map colored by association with schizophrenia in **Fig. 2a**; the top 50 drugs with identified ATC codes were mapped onto another map in **Fig. 2b** (associated data in **Tables S7 and S8 in Supplement 2**). Out of the 13,572 biological pathways, 28 reach Bonferroni significance and 112 have FDR q-value < 5%. For drugs with ATC code, only five are Bonferroni-significant, and 13 have FDR q-value < 5%. Among enriched biological pathways, we find genic intolerance, mental disorders, synapse and neuron pathways, pathways related to histones and nucleosomes, transmembrane transport and ion channels, and epilepsy pathways (**Fig. 2a**). Top drugs on the map are mainly driven by calcium voltage-gated receptors, *DRD2*, acetylcholine receptors, GABA receptors, HCN channels, or some other individual genes (Bonferroni-significant: *DPYD*, *DPP4*, *CCHCR1*, *PSORS1C2*, *CYP17A1*, *MPL*, *NEU1*, *MPL*, *TNF*, *HLA-DQB1*, *ABCB1*). The top FDR-significant drugs targeting calcium channels are cinnarizine, nilvadipine, paramethadione, clevidipine, isradipine, mibefradil, drotaverine, nisoldipine, verapamil, nicardipine and nimodipine.

**Figure 2.**
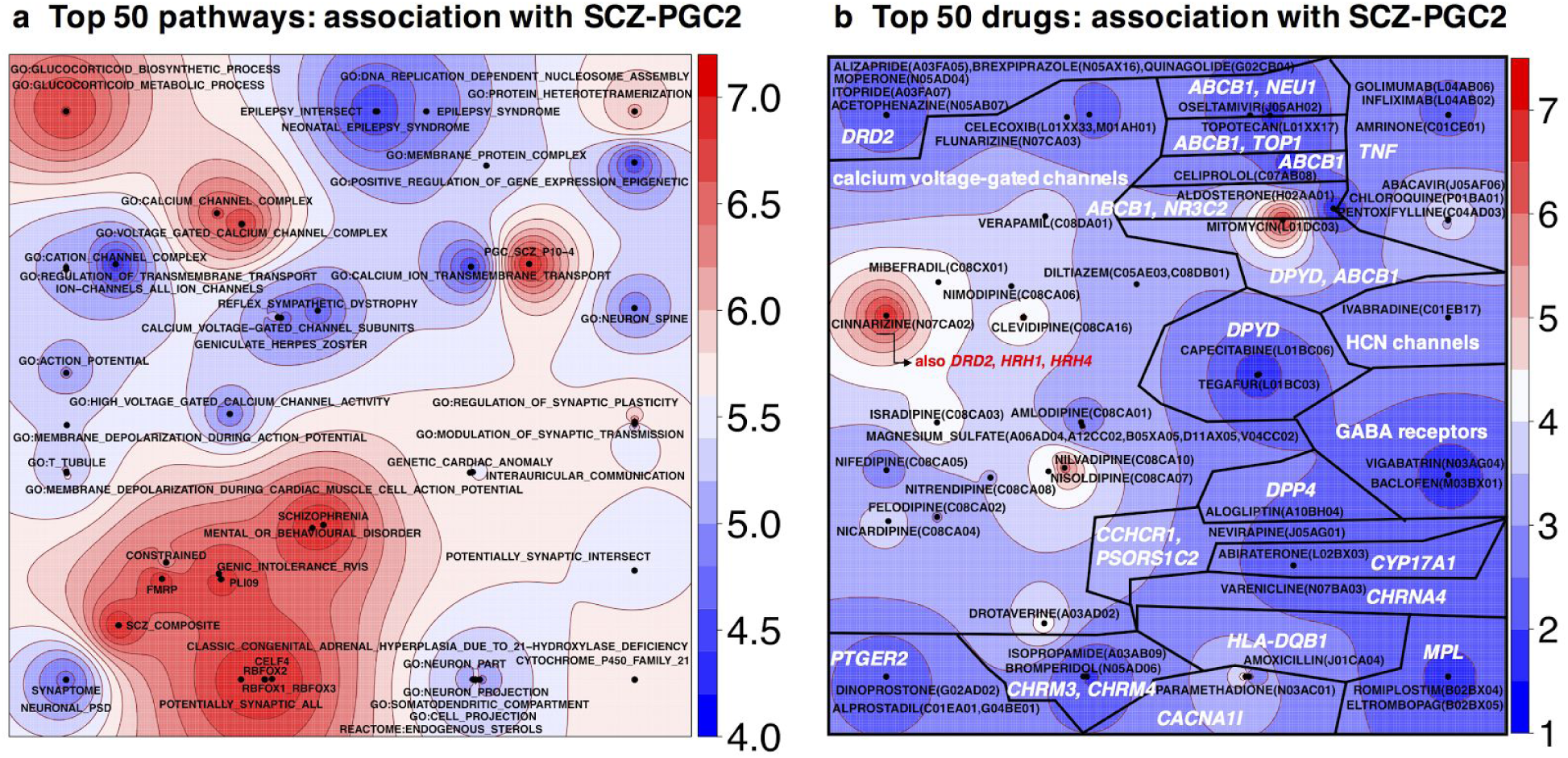
Pathway maps obtained using k-GTM (kernel Generative Topographic Mapping), a dimension reduction algorithm which projects pathways onto a 2D map. The points are gene-sets, positioned according to gene composition. The map is colored by -log_10_(*p*), which measures the degree of association of a gene-set with schizophrenia. (**a**) Top 50 pathways in schizophrenia SCZ-PGC2 GWAS: GO ontology, canonical pathways, gene families or disease gene-sets defined in the Open Targets Platform. All these pathways are FDR-significant according to Benjamini and Hochberg’s q-values, whereas only 28 are Bonferroni-significant. (**b**) Top 50 drugs with identified ATC codes, with target information mined from DGIdb and *K*_i_ DB. Labels indicate the ATC code for each drug (N05 = psycholeptics, etc) as well as the most significant gene(s) in each segment of the map.

### Drug classes enrichment

The enrichment of ATC (Anatomical Therapeutic Chemical) drug classes in the latest schizophrenia GWAS (SCZ-PGC2) is reported in **Fig. 3a**. The enrichment is assessed using the AUC, where AUC = 100% indicates optimal enrichment and AUC = 50% a random result. AUC p-values were computed using Wilcoxon-Mann-Whitney’s test and a Bonferroni threshold (1.10^-3^) was applied to identify significant drug classes, accounting for 49 tests. Antipsychotics (AUC = 75%, *p* = 1.342×10^-9^), selective calcium channel blockers with mainly vascular effects (AUC = 93%, *p* = 3.427×10^-8^), and antiepileptics (AUC = 76%, *p* = 1.814×10^-6^) were significant.

**Figure 3.**
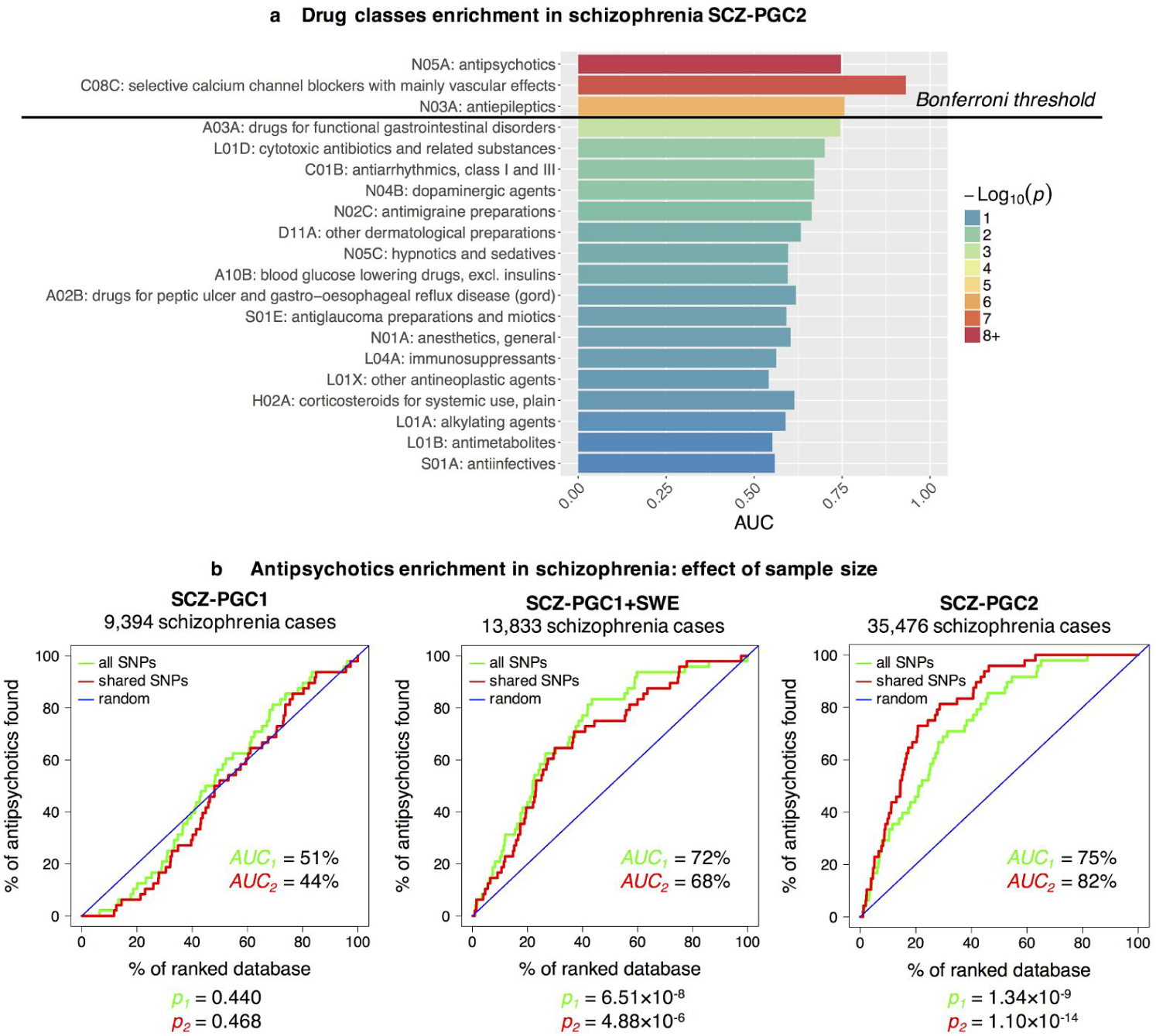
**(a)** Enrichment of top ATC drug classes in SCZ-PGC2 GWAS. AUC is the area under the enrichment curve, and p-values are derived from Wilcoxon-Mann-Whitney’s test, which assesses whether drugs of a given class have a higher association with schizophrenia than expected by chance. **(b)** Antipsychotic enrichment in schizophrenia GWASs as a function of sample size. The figure shows enrichment curves for antipsychotics (ATC code N05A), using three GWASs with increasing sample sizes. The expected “random” enrichment curve is indicated in blue. The red enrichment curve is based on SNPs shared between the three studies, and the green enrichment curve uses all SNPs available in a study. Corresponding areas under the curve (AUC) and p-values (*p*) are provided.

### Effect of sample size on drug class enrichment

Antipsychotics enrichment curves were generated for SCZ-PGC1, SCZ-PGC1+SWE and SCZ-PGC2 (**Fig. 3b**), using only SNPs present in all three studies (“shared SNPs”) or all SNPs available in each study. The p-values associated to the AUC were not corrected for multiple testing, since only three planned comparisons were made. For SCZ-PGC1, the antipsychotics enrichment is equal to a random result (with shared SNPs: *AUC* = 44%, *p* = 0.468); the enrichment is moderate for SCZ-PGC1+SWE (68%, *p* = 4.88×10^-6^), and high (82%, *p* = 1.10×10^-14^) for SCZ-PGC2. As the sample size used in schizophrenia GWAS increases (and consequently the statistical power), so does the enrichment for antipsychotics.

### Protein-protein interaction networks

The proteins targeted by drug classes enriched in SCZ-PGC2 were investigated using protein-protein interaction networks (**Fig. 4a**). The analysis revealed that enriched drug classes (selective calcium channel blockers, antiepileptics and antipsychotics) targeted different subnetworks with association with schizophrenia. Known antipsychotics target dopamine, serotonin, adrenergic, and muscarinic acetylcholine receptors. The selective calcium channel blockers mainly target calcium channels, whereas antiepileptics target GABA receptors, glutamate receptors, sodium channels, calcium voltage-gated channels and nicotinic acetylcholine receptors. The top targets for antipsychotics (with highest association with schizophrenia) are *DRD2*, *CHRM4* and *HTR5A*; for antiepileptics, *CACNA1I* and *SCN9A,* and for selective calcium channel blockers, *CACNA1C* and *CACNB2*. The top genes in epilepsy pathways are *AKT3*, *GABBR2*, and *KCNQ2*, and the main target families are GABA receptors, glutamate receptors, potassium channels, sodium channels, and calcium voltage-gated channels (**Fig. 4b**).

**Figure 4.**
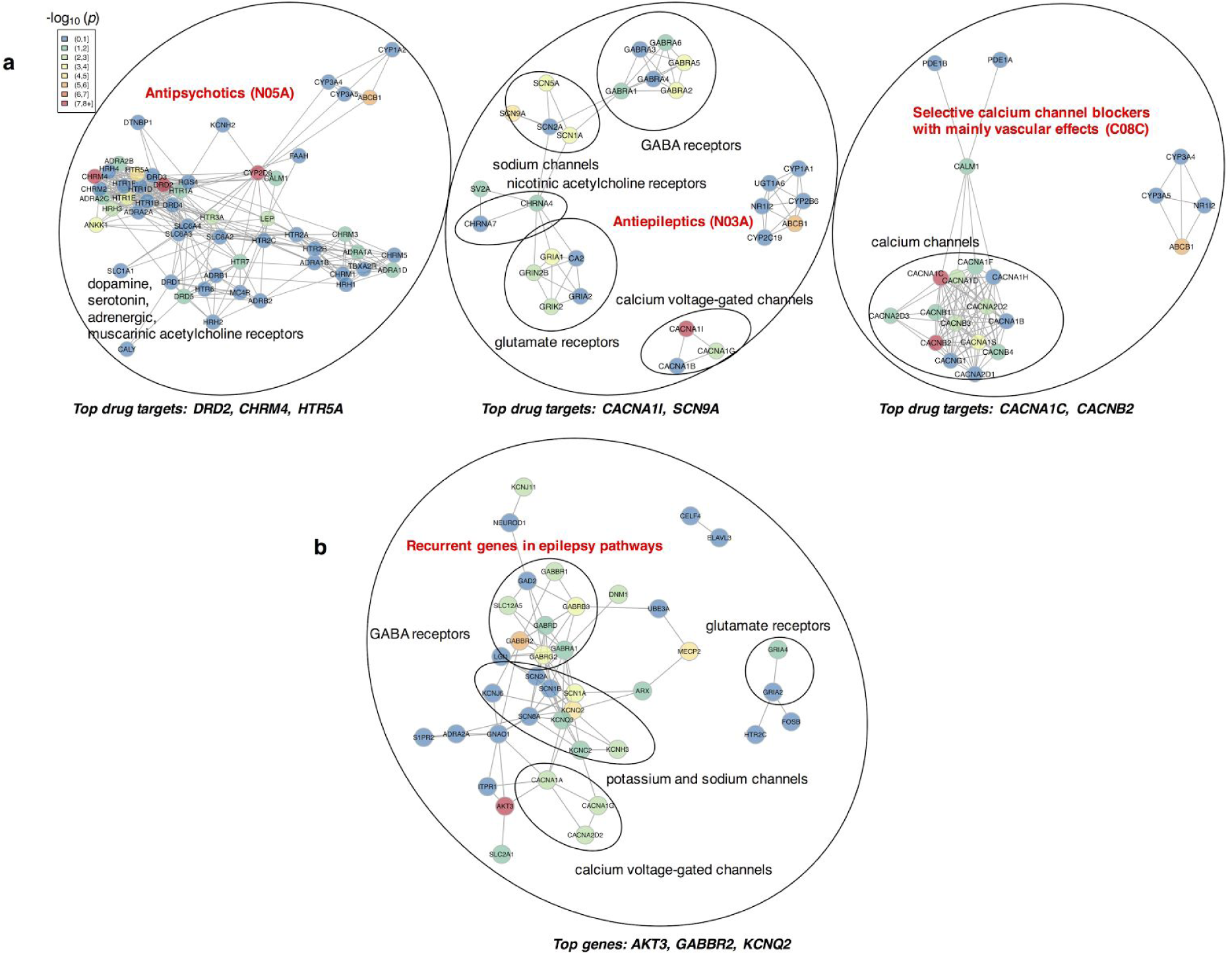
Protein-protein interaction networks. The interactions and interaction scores were obtained through the STRING online platform. Vertices were placed on a plane using the Fruchterman-Reingold layout algorithm. Each node is colored by -log_10_(p), which measures the degree of association of a gene with schizophrenia (SCZ-PGC2). **(a)** Protein-protein interaction networks for the three drug classes significant in SCZ-PGC2 : only proteins targeted by at least 2 drugs within the class are shown. **(b)** Protein-protein interaction network in epilepsy pathways: only genes present in at least 10 epilepsy pathways from Open Targets are shown.

## DISCUSSION

We find that the targets of antipsychotics, the primary drug class used to treat schizophrenia, are enriched for association in current schizophrenia GWAS results. We also show that this enrichment increases with the number of schizophrenia cases included in the GWAS, the largest being SCZ-PGC2 (∼35,000 cases). In addition, our results show significant enrichment for two other drug classes: selective calcium channel blockers and antiepileptics.

It is noteworthy that there is no evidence for a common genetic variant genetic correlation between schizophrenia and epilepsy as measured by linkage disequilibrium score (LDSC) regression ^16^. However, our analyses reveal that epilepsy pathways and the targets of antiepileptics (with GABAergic and antiglutamatergic action) are enriched in schizophrenia. Some antiepileptics have also been investigated for treatment-resistant schizophrenia ^17^.

Voltage-gated channels have been widely studied in psychiatric disorders ^18^, and L-type calcium channels have been associated with schizophrenia in numerous studies ^19^. Amongst top drugs targeting calcium channels, verapamil has been reported to be comparable to lithium for the treatment of mania ^20^. Cinnarizine, which has atypical antipsychotic properties in animal models ^21^, is prescribed for vertigo because of its antihistamine properties and is also an antagonist of dopamine D2 receptors.

We also demonstrated that nicotinic acetylcholine receptors show significant association in SCZ-PGC2. The *CHRNA3-CHRNA5-CHRNB4* gene cluster is strongly associated with schizophrenia; it consists of genes in high LD with each other and has been linked to nicotine dependence ^22^. Some studies indicate that nicotine could have a positive effect on psychotic symptoms and cognitive function in schizophrenic patients ^23^. Our results are consistent with a recent study by Won et al. that also highlights the enrichment of acetylcholine receptor activity in schizophrenia ^24^. Several drugs, such as varenicline and galantamine, target these receptors. Varenicline is a nicotinic agonist used for smoking cessation ^25^ while galantamine is an allosteric modulator of nicotinic receptors and an acetylcholinesterase inhibitor, and has been investigated for the treatment of cognitive impairment in schizophrenia ^26^.

Many of the 162 Bonferroni-significant genes are clustered in the high LD region of the major histocompatibility complex (MHC) and are thus difficult to interpret. However, we also see significance with other individual gene loci encoding calcium voltage-gated channels, sodium channels, potassium channels, cholinergic receptors, and multiple novel genes. The top druggable genes we identify outside the MHC region include *CYP17A1, AS3MT, CACNA1C, NT5C2, DPYD, CACNA1I, XRCC3, FURIN, KCL1, OGFOD2, CKB, CHRNA3, CHRM4, TAOK2, CACNB2, CHRNA5, SRR, CHRNB4, NDUFA4L2, NDUFA13, DRD2.*

The associations with metabolic enzymes such as *CYP2D6*, in which variability may influence antipsychotic plasma levels, is difficult to interpret as the large number of treatment resistant cases included in the PGC2-SCZ GWAS may influence the results; however, recent studies suggest that *CYP2D6* functional variation is not associated with treatment resistance^27^. Compounds targeting proteins encoded by *MCHR1* and *DPP4* might be of particular interest. *MCHR1* antagonists include high affinity ligands such as ATC0175 or ATC0065, which exhibit antidepressive and anxiolytic effects in mouse and rat behavioral models ^28^. *DPP4* inhibitors include gliptins (dutogliptin, alogliptin, etc.) that are used to treat type 2 diabetes, and atorvastatin, which is prescribed due to its cholesterol-lowering properties ^29^. Current antipsychotics can induce insulin resistance ^30^, and drugs which do not or would reverse these effects would be a welcome addition to the pharmacopoeia.

In summary, our workflow may be used to validate the power of a given GWAS, identify new drug targets, and visualise biological pathways. It is suitable for use as a filtering process in the first stages of drug discovery, with detailed target qualification analyses and validation experiments necessary for individual genes and molecules. We conclude that sufficiently powerful GWASs can be examined with increased confidence for drug target identification and repurposing opportunities across complex disorders, by investigating biological pathways, drug gene-sets and druggable genes. In disorders that have few known drug treatments, such as eating disorders and autism, our validation step may be impossible but, once well-powered GWASs with multiple significant signals become available, this approach could still be effective to generate much needed therapeutic hypotheses.

## METHODS

### Methods: Pathway analysis

The pathway analysis software MAGMA v. 1.06 ^31^ was used to generate p-values for genes and gene-sets representing drugs, gene families, biological pathways and disease pathways. GWAS summary statistics are available as SNP p-values, which MAGMA combines to produce gene and gene-set p-values. We used a combined model with top and mean SNP associations to compute gene p-values. These gene p-values are converted to Z-values, which are used as the response variable in a regression model, solved using a generalized least squares approach accounting for linkage disequilibrium. Two types of regression analyses can be conducted: self-contained or competitive. The self-contained approach tests whether the pathway is associated with a trait of interest, whereas the competitive approach tests whether genes in the pathway are more strongly associated than genes outside the pathway. The self-contained approach is more powerful, but it is sensitive to the polygenic nature of observed GWAS statistics inflation and may lead to a higher Type I error ^31,32^. Therefore, we used competitive p-values. In MAGMA, the competitive analysis corrects for gene size, density, minor allele count, and takes into account gene-gene correlations^31^. The SNP positions and frequencies were extracted from the European subset of 1000 genomes phase III v.5a ^33^ with genome assembly hg19. We used Ensembl release 75 ^34^ for the gene positions. The gene window was set to 35 kb upstream and 10 kb downstream in MAGMA to include gene regulatory regions. We generated FDR-adjusted p-values or *q-values* for genes and gene-sets, using Benjamini and Hochberg’s method to account for multiple testing ^35^; we also provide the Benjamini and Yekutieli q-values^36^ and Bonferroni-corrected p-values for all our results in **Supplement 2**.

### Methods: Pathway maps

The top 50 biological pathways and top 50 drugs with ATC codes were used to produce two separate maps, using the p-values obtained in the pathway analysis step. Gene-sets were encoded by gene content (binary vectors) and a Tanimoto similarity matrix was generated. This matrix was used as input for the k-GTM algorithm ^14,15^ implemented in GTMapTool v1.0 ^37^. Five parameters need to be defined by the user: the square root of the number of sample points (*k*), the square root of the number of radial basis functions (RBF), the regularization coefficient (*l*), the RBF width factor (*w*) and the feature space dimension (*D*). We set *k* = square root of the number of data points in the input kernel, *m* = square root of *k*, *l* = 1 and *w* = 1 (default values). The feature space dimension *D* was estimated as the number of PCs explaining 99.5% of the variance in the input data. We used the same method to compute the number of independent tests and generate the Bonferroni-corrected p-values for pathways. The maps were colored by schizophrenia association in -log_10_(*p*) units using the kriging algorithm implemented in the R package *gstat* ^38^.

### Methods: Protein-protein interaction networks

Genes driving the association in pathway clusters or drug families were highlighted in protein-protein interaction networks. Protein-protein interaction scores were generated using the STRING v.10 online platform ^39^, which integrates information from genomic context predictions, high-throughput lab experiments, co-expression, automated textmining, and other databases. The Fruchterman-Reingold layout algorithm implemented in the R package *igraph* ^40^ *was used to position the vertices on the graphs depending on the interaction score; each gene (node) was colored by its association with schizophrenia computed by MAGMA, in –*log_10_(*p*) units.

### Methods: Enrichment measure for groups of gene-sets

Instead of investigating individual gene-sets, we focused on *groups* of gene-sets. For example, a class of drugs can be represented by a group **G** of drugs (gene-sets). To determine whether **G** is significantly enriched, we can draw enrichment curves, widely used in virtual screening ^41^. The curves display the percentage of hits found when decreasing the value of a scoring function. Here, the scoring function is the gene-set association with the trait of interest in –log_10_(*p*) units, and the hits are the gene-sets in **G**. The area under this enrichment curve (AUC) provides a quantitative assessment of the enrichment of **G** in a GWAS and is computed using the trapezoidal approximation of an integral. The expected random result is *AUC* = 50% and the maximum value is *AUC* = 100%.

The AUC significance was assessed using Wilcoxon-Mann-Whitney test (WMW), which tests whether the data distribution is the same within two different groups (e.g., gene-sets in **G** and not in **G**) ^42^ - also, the AUC can be directly calculated from the Wilcoxon-Mann-Whitney U statistic ^43^. We used this enrichment measure to assess whether drugs in a set **G** were more associated with a disorder than others, while accounting for the fact that drug gene-sets are diverse and noisy, due to an incomplete knowledge of targets, the presence of off-targets without any association with the disorder, and the fact that drugs may have different mechanisms of action within the same therapeutic class.

### Materials: Schizophrenia GWAS summary statistics

In this paper, we used three GWASs conducted in 2011 ^12^, 2013 ^13^ and 2014 ^4^ with increasing sample sizes (cf. **Fig. 3b** and **Table S1 in Supplement 1**). The three studies were coined SCZ-PGC1, SCZ-PGC1+SWE and SCZ-PGC2, respectively. The three studies mainly contain individuals of European ancestry ^4,12,13^; SCZ-PGC2 is the only study including the X chromosome and individuals with East Asian ancestry. Only SNPs present in the European subset of 1000 genomes phase 3 v.5a ^33^ with minor allele frequency (MAF) ≥ 1% were kept. The genomic inflation factor as well as the LD score intercept were computed for each set using the LDSC software v. 1.0.0 ^44^. All p-values were subsequently corrected using the LD score intercept - a score based on linkage disequilibrium that should provide a better way to control for inflation than the genomic inflation factor ^45^. Only the 1,123,234 SNPs shared among SCZ-PGC1, SCZ-PGC1+SWE and SCZ-PGC2 were considered when comparing the three studies. The latest and most powerful GWAS (SCZ-PGC2) was used to investigate the enrichment of drug classes, drug gene-sets, and biological pathways.

### Materials: Drug gene-sets

Drug/gene interactions are mainly derived from drug/target activity profiles. The data was drawn from two sources: the Drug-Gene Interaction database DGIdb v.2 ^46^, and the Psychoactive Drug Screening Database *K*_i_ DB ^47^ downloaded in June 2016. DGIdb is a new resource that integrates drug/gene interactions from 15 databases, amongst which DrugBank and ChEMBL; the data is directly available as drug/gene pairs and genes are identified by their HGNC (HUGO Gene Nomenclature Committee) names ^48^. *K*_i_ DB provides *K*_i_ values for drug/target pairs and is particularly relevant for psychoactive drugs; only human assays were considered and drug/target pairs were kept if 5 < p*K*_i_ < 14. More details on the filtering procedure can be found in **Text S1 in Supplement 1**. Gene-sets were produced by merging both DGIdb and *K*_i_ DB drug/gene data and by converting HGNC names to Ensembl release 75 ^34^ identifiers. The number of unique gene-sets was 3,939 at the end of the filtering process, 3913 with variants in SCZ-PGC2 (2586 independent gene-sets), out of which 1026 were mapped to at least one ATC code. We annotated groups of drugs using ATC classes, listed in **Table S9 in Supplement 2** and containing at least 10 drugs. The “validation” set for schizophrenia GWASs is the set of antipsychotics with ATC code N05A - all schizophrenia drugs belong to this class (cf. **Table S2 in Supplement 1** for the list of prescription drugs in the UK).

### Materials: Biological pathways

We refer to our ensemble of gene ontology pathways, canonical pathways, disease pathways, and gene families as “biological pathways”. Canonical (CP) and Gene Ontology (GO) gene-sets were extracted from MSigDB v5.2^49^. MSigDB is a regularly updated resource gathering pathways and ontologies from the main online databases. CP sets were curated from: BioCarta, KEGG, Matrisome, Pathway Interaction Database, Reactome, Sigma Aldrich, Signaling Gateway, Signal Transduction KE and SuperArray. These “pathways” provide a practical way to investigate the function of a subnetwork without accounting for the complexity of biological networks. Disease pathways were extracted from the Open Targets platform ^50^ in January 2017 and gene families were identified using information provided on the HGNC website. The total number of biological pathways was 13,572 (9408 independent pathways).

## Supplementary Materials

**Supplement 1: PDF File:**

Supplementary Figures 1-2, Supplementary Tables 1–5, and Supplementary Texts 1-2

**Supplement 2: Excel File:**

Supplementary Tables 6-10

## Acknowledgments

*K*_i_ determinations were generously provided by the National Institute of Mental Health’s Psychoactive Drug Screening Program, Contract # HHSN-271-2013-00017-C (NIMH PDSP). The NIMH PDSP is Directed by Bryan L. Roth MD, PhD at the University of North Carolina at Chapel Hill and Project Officer Jamie Driscoll at NIMH, Bethesda MD, USA.

## Author contributions

H.A.G. produced the results and introduced the pathway visualisation methodology, G.B. provided advice and guidelines, both H.A.G. and G.B. contributed to the methodology and the writing of the paper.

## Additional information

### Data and materials availability

All data used in this paper are freely available online (cf. references and supplementary materials).

